# Deciphering the mystery of influenza virus A antigenic drift and immune escape

**DOI:** 10.1101/2020.12.23.424101

**Authors:** Yongping Ma, Changyin Fang

## Abstract

The biggest challenge of influenza A virus vaccine development is the constant mutation of the virus, especially the antigenic drift. Therefore, we suggest establishing a research system for upgrading of current influenza A vaccines. The most fundamental but neglected thing is to determine the immune escape mutation (IEM) map (IEMM) of 20 amino acids in linear epitope to reveal the precise mechanism of the viral immune escape. Based on this assumption, we primarily divided IEM of HA-tag linear epitope into four types according to antibody affinity by ELISA. Our aim is to attract scholars to establish of IEMM library to deal with influenza A virus mutations in the future.

## Introduction

The crucial factor of the annually prevalence of seasonal human influenza A virus is the evolution of the viruses to escape the immunity induced by prior infection or vaccination (*1*). That is known as antigenic drift and reassortment (*2*). We will be interested only in antigenic drift in this study. Antigenic drift results in considerable mismatches between current vaccines and circulating strains (*3–5*). The dilemma is that the vaccine development cannot keep pace with viral mutations. Therefore, current seasonal influenza vaccines are ineffective against pandemic influenza and cause global morbidity and mortality annually. Even though the human seasonal influenza vaccine strains are optimized according to the genome-sequencing data of antigenic drift, mismatches often occur and lead to a decreasing in vaccine effectiveness (*5, 6, 7*). Therefore, the vaccine strains must be updated and re-administered annually. Normally, it takes about 6 months to manufacture matched vaccines for the circulating viruses (*8*). Therefore, predicting the next season’s predominant influenza strains is a major job of WHO. However, that is often incorrect (*2, 5, 9–11*). Thus, development of universal influenza virus vaccines is of great concern recent decades (*8, 11*). The potential side effects are antibody-dependent enhancement (ADE) disease by virus fusion-enhancing for mismatched vaccines (*12, 13*).

Antigenic drift is the typical process of human influenza virus evolution by natural selection of mutations in the hemagglutinin (HA) that enable the virus to evade the pre-existing human immune response (*14*). According to antigenic drift successfully model, new strains may be generated constantly through mutation, but most of these cannot expand in the host population due to pre-existing immune responses against epitopes of limited variability (*1*5). Obviously, this natural selection is biased, *e.g.*, the amino acid at position 147 of human H1N1 is mutated from R (Arg, 1977-liker) to K (Lys,1991-liker, 2009-liker) as escape mutants, not random occurrence of 20 amino acids *(16)*. However, the exact mechanisms of its natural selection is unknown that involves in the interactions between the prior antibodies and the mutated HA. We consider the essence of virus antigenic drift is the change of the degree of affinity between the novel residues in antigenic epitope and pre-existing antibodies. Therefore, it is very important to determine the laws of amino acid mutation and the degree of affinity to pre-existing antibodies for vaccine design and virulence prediction of epidemic strains. Therefore, the first thing is to uncover the secret between key amino acid mutations and immune escape. Given the complexity of antigen spatial epitopes, this study just focuses on the linear epitope mutation and immune escape.

Here, the enzyme linked immunosorbent assay (ELISA) was employed to reveal the precise mechanism of the viral immune escape by measuring the degree of affinity between anti-HA-tag (Y1P2Y3D4V5P6D7Y8A9) monoclonal antibodies (mAbs) and HA-tag mutants.

In order to describe the relationship between linear epitope mutation and immune escape, we cautiously introduced four concepts: (i) **Immune escape mutation (IEM)** means that the substituted residue causes the antigen to lose its affinity (recognition) to pre-existing antibody, or to remain less than assumed 30% affinity without neutralization. (ii) **ADE (antibody-dependent enhancement) risk mutation (ADERM)** refers to that the substituted residue causes the antigen to remain pre-existing antibody affinity more than assumed 30% but less than 50%. However, pre-existing antibody could not neutralize the mutated antigen/pathogen (*17*). Rather, the virus-Ab complex with low affinity enhanced virus uptake resulting from the attachment of immune complexes to Fcγ receptor and enhanced the infection (*17, 18*). (iii) **Equivalent mutation (EQM)** means that the substituted residue causes the antigen to remain pre-existing antibody affinity beyond the assumed 60 % but less than 80 %. Fortunately, the pre-existing antibody still completely neutralizes the antigen. (iv) **Invalid mutation (IVM)** means the substituted residue does not affect the pre-existing antibody affinity and the antibody completely neutralizes the antigen.

## Results

### Mapping of key residues in HA-tag to different mAbs

HA-tag (Y1P2Y3D4V5P6D7Y8A9) was substituted one-by-one from Y1 to A9 with G, E or H, respectively. Except A9 residue, mAb TA180128 (Origene, USA) reacted to all of the residues in HA-tag. Y8G, D7G, Y8E, D7H, Y8H, V5I, Y3H, V5T, V5A, V5L, V5S, V5M, Y1H, Y1E, and V5K mutants were IVM for maintaining more than 80% affinity to mAb TA180128. P2H, D4E, V5F, P2E, P6G, and D4H mutants were kept less than 8% affinity, and P2G, P6H, V5Y, V5E, V5D, V5W, P6E, P6A, P6T, P6K, P6Y, P6S, P6C, P6V, P6L, P6M, P6I, P6W, P6F, P6Q, P6D, P6R, and P6N mutants were lost the affinity to mAb TA180128 completely as IEM (Figure 1). Meanwhile, Y1G, Y3G, V5G and Y3E maintained about 16-18% affinity as IEM, therefore, this 33 mutants were IEM to mAb TA180128. Meanwhile, V5R, V5C and V5N maintained about 35%, 41% and 48% affinity, respectively and were ADERM. V5H, V5P, V5Q, D7E, D4G, and V5P were EQM and maintained about 66%-79% affinity to mAb TA180128, respectively (Figure 1). However, mAb 4ab000002 (4A Biotech, Beijing) only reacted to the P6, D7 and Y8 residues in HA-tag. P6I maintained 42% affinity as an ADERM. D7H maintained less than 10% affinity, and P6A, P6Y, Y8G, P6C, P6Q, P6N, P6K ,P6D, P6R, P6W, P6F, P6H, and Y8E mutants were lost the affinity completely as an IEM (Figure 2). However, D7G, D7E, P2E, D4E, A9E, P6E,V5E, and P6M mutants were EQM for maintained more than 62% to 79% affinity (Figure 2). P6T, P6S, P6V, V5D, V5I, V5P,V5Y, V5Q, V5C, V5F, V5N, V5T, V5W, V5M, Y8H, V5S,V5K,V5L,V5A, V5H,Y3G, V6R, A9H, Y1H, P2H, Y3H, D4H, P6G, V5G, D4G, A9G, Y3E, P2G, and Y1G mutants maintained more than 80% affinity and were IVM to mAb 4ab000002 (Figure 2).

**Figure 1.**
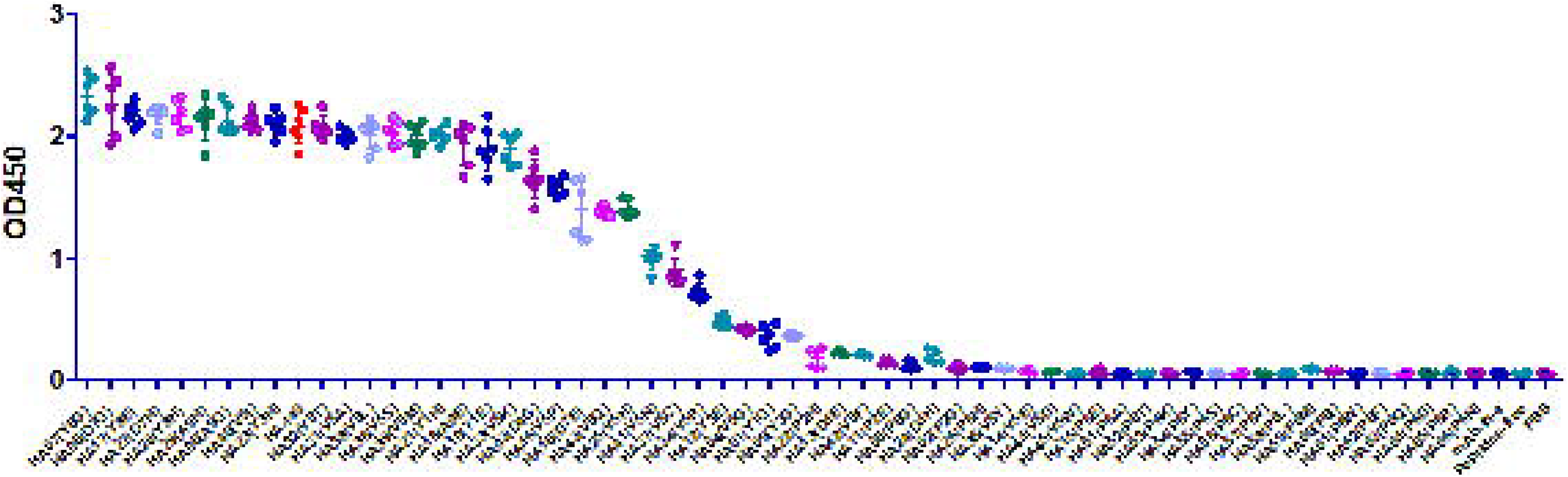
Scanning the HA-tag mutants-mAb TA180128 affinity.

**Figure 2.**
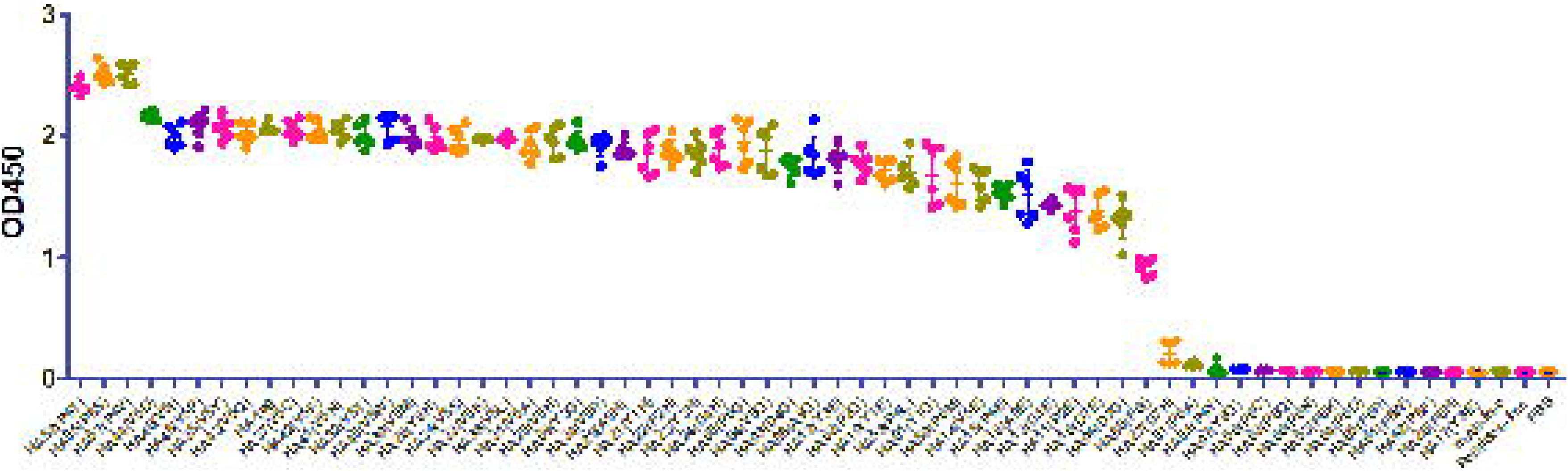
Scanning the HA-tag mutants-mAb 4ab000002 affinity.

### A half of amino acids substitution was IVM and EQM

Objective to reveal the law of IEM, V5 and P6 of HA-tag were totally substituted with the other 16 amino acids, respectively. As a summary, V5 was substituted by F and G maintained less than 16% affinity and by Y, E, D and W were lost affinity as IEM accounted for 31.6% of the 19 amino acids. Otherwise, V5 was substituted by H, Q, and P were EQM accounted for 15.8% and maintained about 66%-79% affinity to mAb TA180128, respectively (Figure 1). That was, more than 47% substitutions were completely IVM and EQM. However, V5 was substituted by I, T, A, L, S, M and K, as IVM, were remained more than 80% affinity to mAb TA180128 accounted for 36.8%. Meanwhile, V5 was substituted by R, C and N maintained about 35%, 41% and 47.8% affinity, respectively and were ADERM accounted for 15%.

Similarly, P6 was substituted by I maintained 42% affinity as an ADERM. P6 was substituted by A, Y, C, Q, N, K, D, R, W, F and H were lost the affinity completely as an IEM accounted for 57.8%. However, P6 was substituted by E and M were EQM for maintained more than 71% affinity accounted for 10.5%. P6 was substituted by T, S, V, L and G maintained more than 80% affinity accounted for 26.3% and were IVM to mAb 4ab000002 (Figure 2). That was, more than 36.8% substitutions were completely IVM and EQM.

### The aryl group in amino acid side chains was more destructiveness

Due to the lack of X-ray or Cryo-EM data, it was difficult to explain the mechanisms of P6 residue IEM. However, there were two conclusions: (1) both the V5 and P6, substituted by aryl amino acids (F, Y, W), were more destructive, leading to IEM. (2) Key amino acid substituted by the same polarity residue did not guarantee the affinity to mAb unchanged (Figure 3a, b).

**Figure 3.**
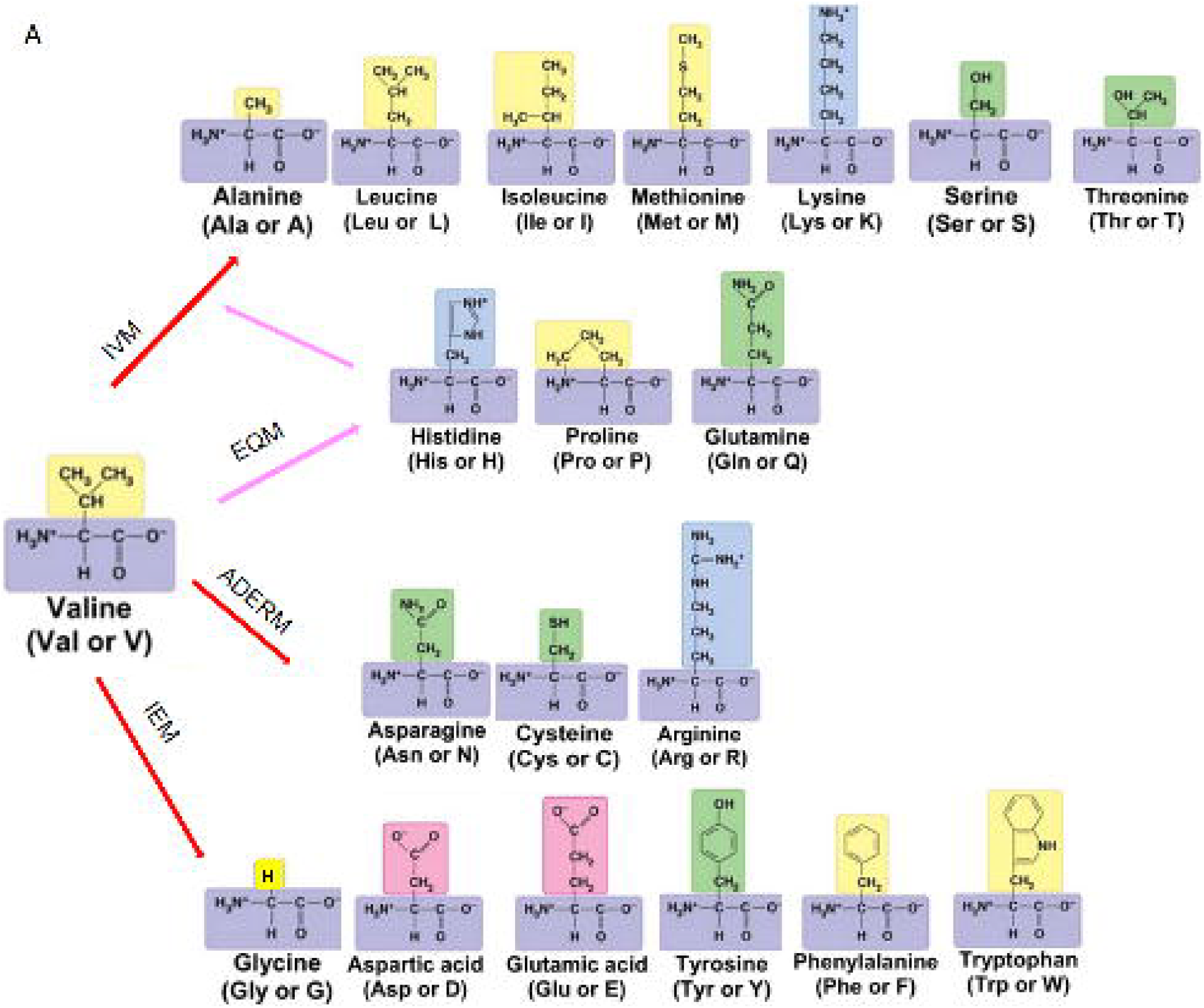
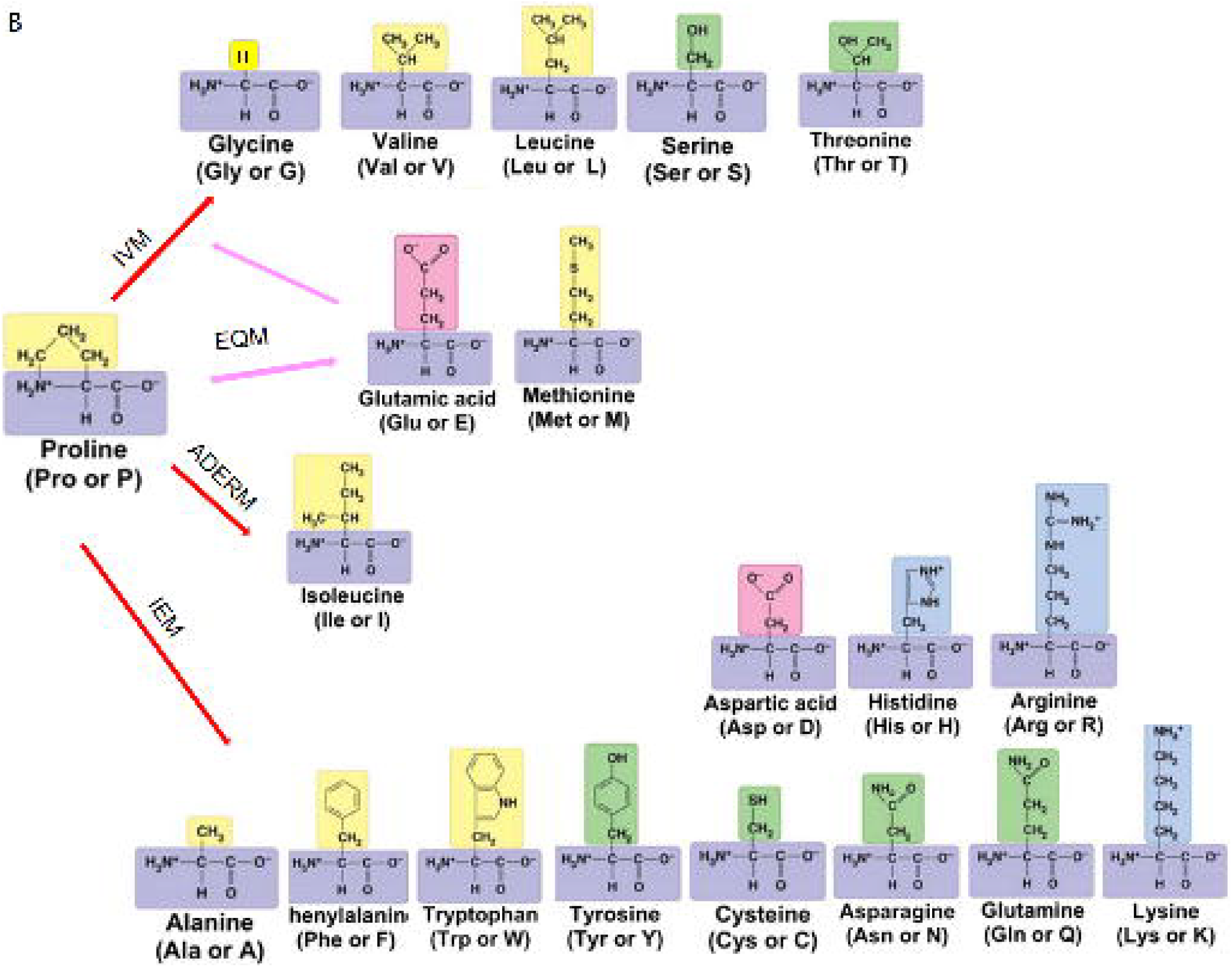
Schematic of IEMs. (A) Schematic of V5 of HA-tag substituted by the other 19 amino acids. V5G, V5D,V5E, V5F, V5Y and V5W were IEMs. (B) Schematic of P6 of HA-tag substituted by the other 19 amino acids. P6A, P6C, P6D, P6H, P6K, P6N, P6Q, P6R, P6F, P6Y and P6W were IEMs.

## Discussion

The process of antigenic drift of human influenza A virus occurs constantly during the replication cycle for without proofreading mechanisms (*19*). Theoretically, the substitution of key amino acid residue occurs in human influenza A virus by the other amino acids randomly. Actually, a great deal of mutations did not exist in nature for immune pressure (*15*). The theory suggested that most of the constantly generated new strains could not expand in the host population due to pre-exist immune responses (*15*). So far, it could not predict which kinds of amino acid substituted to a particular amino acid residue in specific epitope leading to immune escape. Thus, it was a significant work to determine the immune escape mutation map **(IEMM)** of 20 amino acids.

IEMM required the help of Cryo-EM and X-ray assay that was known a time-consuming and costly item. For lack of 3D structure data, our research, serving as a starting point for this novel field of IEMM, was a little superficial.

However, we believed that IEMM was a virgin land of immunology development in the future. Our research suggested that a half of amino acid mutation was IVM in a specific position. Therefore, epidemiologists could judge the infectivity and/or antigenicity of a new mutant strain based on the rule of IEMM and focus on ADERM and IEM to stop or decrease ADE and immune escape (*17*). The vaccinologists just focused on the IEM and ADERM, designed different epitope mutants as epitope library and determined their antigenicity and safety. In the event of IEM or ADERM strain pandemic, the matching epitope library sequences would be immediately constructed into pre-existing vector to substituted of the original (parental) sequence or added at the C-terminus for purified subunit vaccine and/or recombinant live vaccine develop (Figure 4). For severe infectious diseases such as H7N9 and SARS-COV-2, vaccine manufacturers could develop mAb library according to IEM or ADERM linear epitope library in advance. In case an IEM strain pandemic, the matching mAb in library would be immediately produced for emergency usage. In this way, the strategy for seasonal influenza virus prevention would be changed from chase to ambush.

**Figure 4.**
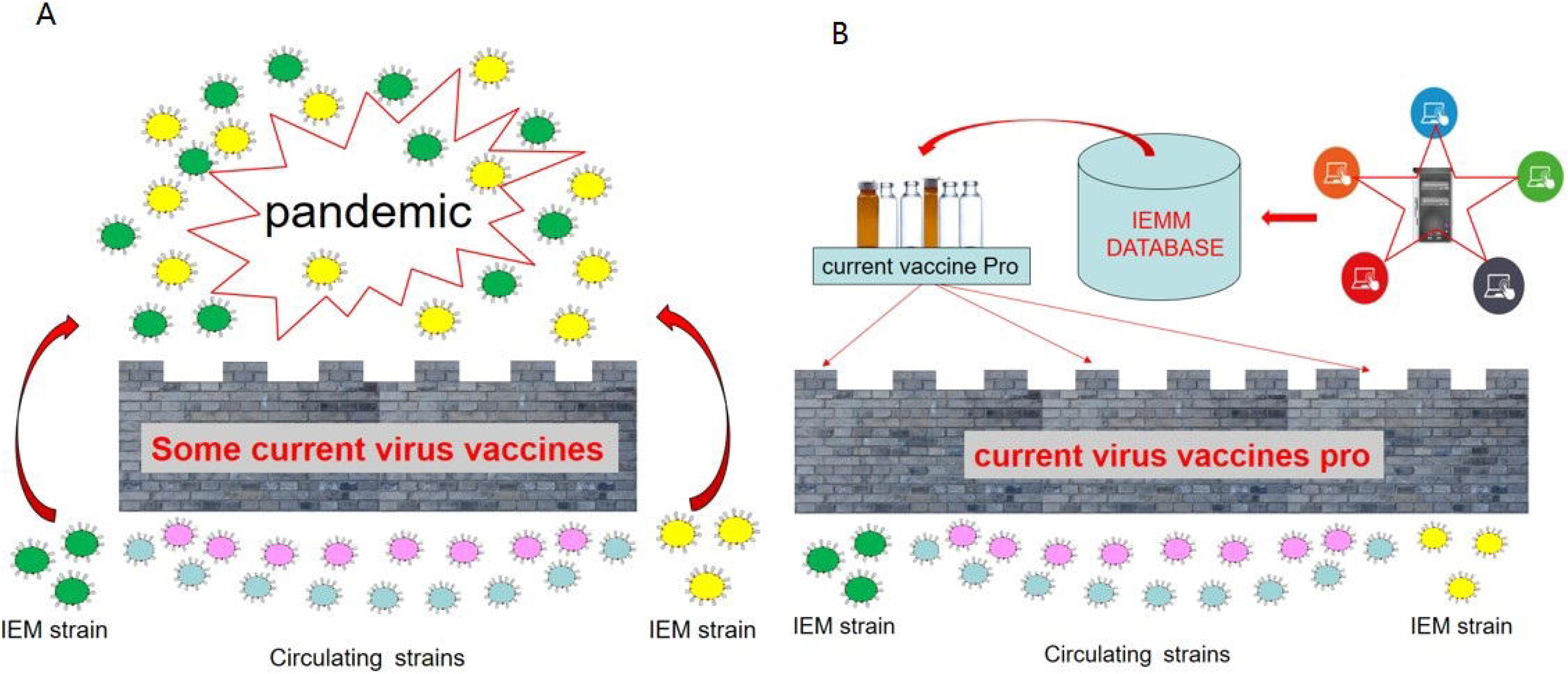
IEMM vision and future vaccine improvements. (A) Antigenic drift caused circulating virus strains to mismatch with current vaccine strains, leading to IEM and pandemic. (B) Based on IEMM database, current vaccine strains were improved to match with circulating virus strains (antigenic drift strains).

Theoretically, as a common sense, antigenic residue determines antibody’s structure. However, it is unknown that the specific amino acid X on the epitope interacts preferentially with the amino acids list on the antibody. Furthermore, what kinds of substitution at critical binding sites could significantly destroy the binding affinity between antigen-antibody and subsequently cause immune escape.

Moreover, it had great theoretical significance to investigate the effects of amino acids next to N- and C-terminus of the mutated residue on IEMM. For example, whether the IEMM of D4 and D7 of HA-tag were inconsistent due to the difference residues next to them was worth studying in future. However, due to the lack of dataset, it is not possible to evaluate quantitatively the four mutations (IEM, EVM, ADERM and IVM) mentioned above and their effects on virus-receptor binding.

## Materials and Methods

### Peptides and agents

Peptides of HA-tag and its mutants (more than 98% purity) were synthesized commercially by Nanjing Yuanpeptide Biotech (Nanjing, China) (Table S1). Mouse anti-HA-tag mAb were purchased from Qrigene (TA180128, MD, USA) and 4A Biotech (4ab000002, Beijing, China), respectively. HRP conjugated AffiniPure goat anti-mouse IgG (Proteintech, SA00001-15, Wuhan, , China) and DAB color development kit (AR1026) were purchased from Biological company. The other buffers were prepared in our laboratory.

### ELISA

The peptides of HA-tag and its mutants were diluted to 2.0 μg/ml and each was added 50 μl into 96 well plate for 1 h to adsorb (n=3). After washing with phosphate buffer saline containing 0.05% Tween-20 (PBST) three times, the plate was blocked with rapid blocking buffer for 10 min. Then, washing with PBST three times, mouse anti-HA-tag lgG2a mAb (4A.Biotech, 4ab000002, Beijing, China 1:1000) or lgG1 mAb (Origene, TA180128, USA, 1:2000) were added into each well (40 μl/well), respectively, and incubated at 37℃ for 1 h. After washing and blocking as previous described, 100 μl of horseradish peroxidase (HRP)-labeled goat anti-mice IgG (Proteintech, SA00001-15, Wuhan, China, 1:2500) were added and incubated at 37℃ for 1 h. After color developing with TMB single-component substrate solution (Solarbio, PR1200, Beijing, China) and stopping, the results were detected by plate reader, respectively (*20*). Peptides-free and mAb-free wells were performed as negative controls. The OD450 was valued for analysis.

## Supporting information

Supplemental Ttable 1

## Acknowledgements

No

## Contributions

Y.M. conceived of the paper and written the manuscript; C.F. contributed, as co-first author, to perform the experiments, analyze data and prepare figures.

## Supplementary material

Supplementary table 1. Sequences of HA-tag and its mutants.

