## Supplemental Ttable 1 for "Deciphering the mystery of influenza virus A antigenic drift and immune escape"

**Table S1**	Sequences of HA-tag and its mutants

| Characteristics | Sequence |
| --- | --- |
| HA-tag | PTRSEWECRCSGSSYPYDVPDYA |
| HA01（Y1G） | PTRSEWECRCSGSS**G**PYDVPDYA |
| HA02（P2G） | PTRSEWECRCSGSSY**G**YDVPDYA |
| HA03（D4G） | PTRSEWECRCSGSSYPY**G**VPDYA |
| HA04（V5G） | PTRSEWECRCSGSSYPYD**G**PDYA |
| HA05（P6G） | PTRSEWECRCSGSSYPYDV**G**DYA |
| HA06（D7G） | PTRSEWECRCSGSSYPYDVP**G**YA |
| HA07（Y8G） | PTRSEWECRCSGSSYPYDVPD**G**A |
| HA08（A9G） | PTRSEWECRCSGSSYPYDVPDY**G** |
| HA09（Y3G） | PTRSGWECRCSGSSYP**G**DVPDYA |
| HA10（Y1E） | PTRSAWECRCSGSS**E**PYDVPDYA |
| HA11（P2E） | PTRSTWECRCSGSSY**E**YDVPDYA |
| HA12（Y3E） | PTRSKWECRCSGSSYP**E**DVPDYA |
| HA13（D4E） | PTRSRWECRCSGSSYPY**E**VPDYA |
| HA14（V5E） | PTRSYWECRCSGSSYPYD**E**PDYA |
| HA15（P6E） | PTRSSWECRCSGSSYPYDV**E**DYA |
| HA16（D7E） | PTRSCWECRCSGSSYPYDVP**E**YA |
| HA17（Y8E） | PTRSVWECRCSGSSYPYDVPD**E**A |
| HA18（A9E） | PTRSLWECRCSGSSYPYDVPDY**E** |
| HA21（Y1H） | PTRSAWECRCSGSS**H**PYDVPDYA |
| HA22（P2H） | PTRSTWECRCSGSSY**H**YDVPDYA |
| HA23（Y3H） | PTRSKWECRCSGSSYP**H**DVPDYA |
| HA24（D4H） | PTRSRWECRCSGSSYPY**H**VPDYA |
| HA25（V5H） | PTRSYWECRCSGSSYPYD**H**PDYA |
| HA26（P6H） | PTRSSWECRCSGSSYPYDV**H**DYA |
| HA27（D7H） | PTRSCWECRCSGSSYPYDVP**H**YA |
| HA28（Y8H） | PTRSVWECRCSGSSYPYDVPD**H**A |
| HA29（A9H） | PTRSLWECRCSGSSYPYDVPDY**H** |
| HA31(V5A) | PTRSEWECHCSGSSYPYD**A**PDYA |
| HA32(V5T) | PTRSEWECACSGSSYPYD**T**PDYA |
| HA33(V5K) | PTRSEWECTCSGSSYPYD**K**PDYA |
| HA34(V5R) | PTRSEWECKCSGSSYPYD**R**PDYA |
| HA35(V5Y) | PTRSEWECYCSGSSYPYD**Y**PDYA |
| HA36(V5S) | PTRSEWECSCSGSSYPYD**S**PDYA |
| HA37(V5C) | PTRSEWECCCSGSSYPYD**C**PDYA |
| HA38(V5L) | PTRSEWECVCSGSSYPYD**L**PDYA |
| HA39(V5M) | PTRSEWECLCSGSSYPYD**M**PDYA |
| HA40(V5I) | PTRSEWECMCSGSSYPYD**I**PDYA |
| HA41(V5N) | PTRSEWECICSGSSYPYD**N**PDYA |
| HA42(V5F) | PTRSEWECNCSGSSYPYD**F**PDYA |
| HA43(V5Q) | PTRSEWECFCSGSSYPYD**Q**PDYA |
| HA44(V5D) | PTRSEWECQCSGSSYPYD**D**PDYA |
| HA45(V5W) | PTRSEWECDCSGSSYPYD**W**PDYA |
| HA46(V5P) | PTRSEWECWCSGSSYPYD**P**PDYA |
| HA51(P6A) | FGEVFNATRFASVYAWNRKRIYPYDV**A**DYA |
| HA52(P6T) | SNCVADYSVLYNSASFSTFKCYGVSYPYDV**T**DYA |
| HA53(P6K) | NDLCFTNVYADSFVIRGDEVRQIAYPYDV**K**DYA |
| HA54(P6Y) | GCVIAWNSNNLDSKVGGNYNYPYDV**Y**DYA |
| HA55(P6S) | FNCYFPLQSYGFQPTNGVGYQYPYDV**S**DYA |
| HA56(P6C) | YQPYRVVVLSFELLHAPYPYDV**C**DYA |
| HA57(P6V) | TVEKGIYQTSNFRVQPTESIVRFPYPYDV**V**DYA |
| HA58(P6L) | APGQTGKIADYNYKLPDDFYPYDV**L**DYA |
| HA59(P6M) | NYLYRLFRKSNLKPFERDISTEIYQAYPYDV**M**DYA |
| HA60(P6I) | EIYQAGSTPCNGVEGFNCYYPYDV**I**DYA |
| HA61(P6W) | FVFLVLLPLVSSQCYPYDV**W**DYA |
| HA62(P6F) | HADQLTPTWRVYSTGSNVYPYDV**F**DY |
| HA63(P6Q) | IEDLLFNKVTLADAGFIKYPYDV**Q**DY |
| HA64(P6D) | FKEELDKYFKNHGSGSGYPYDV**D**DYA |
| HA65(P6R) | DISTEIYQAGSTPCNGVEGFNCYYPYDV**R**DY |
| HA66(P6N) | TESNKKFLPFQQFGRDIADTTDAVRDPYPYDV**N**DY |

1. HA-tag was underlined. 2. Substitution residue was showed in bold type.
